# Parvalbumin Neurons and Cortical Coding of Dynamic Stimuli: A Network Model

**DOI:** 10.1101/2022.09.22.509092

**Authors:** Jian Carlo Nocon, Isaac Paul Boyd, Howard Gritton, Xue Han, Kamal Sen

## Abstract

Cortical circuits feature both excitatory and inhibitory cells that underlie the encoding of dynamic sensory stimuli, e.g., speech, music, odors, and natural scenes. While previous studies have shown that inhibition plays an important role in shaping the neural code, how excitatory and inhibitory cells coordinate to enhance encoding of temporally dynamic stimuli is not fully understood. Recent experimental recordings in mouse auditory cortex have shown that optogenetic suppression of parvalbumin neurons results in a decrease of neural discriminability of dynamic stimuli. Here, we present a multilayer model of a cortical circuit that mechanistically explains these results. The model is based on parvalbumin neurons which respond to both stimulus onsets and offsets, as observed experimentally, and incorporates characteristic shortterm synaptic plasticity profiles of excitatory and parvalbumin neurons. We reveal that by tuning the relative strengths of onset and offset inputs to parvalbumin neurons, the model generates different regimes of coding dominated by rapid firing rate modulations or spike timing. Moreover, the model replicates the experimentally observed reduction in neural discrimination performance during optogenetic suppression of parvalbumin neurons. These results suggest that distinct onset and offset inputs to parvalbumin neurons enhance cortical discriminability of dynamic stimuli by encoding distinct temporal features, enhancing temporal coding, and reducing cortical noise.

**New & Noteworthy:** Here we propose a model for the mechanisms that underlie neuron responses in the auditory cortex. This study focuses on single channel tuning of simple interactions between neurons. Using physiologically relevant parameters that underly parvalbumin and excitatory neurons, in the proposed artificial network, we show that we can recreate observed results in live studies.

## Introduction

Temporal processing of dynamic stimuli is critical for human perception of behaviorally relevant stimuli, e.g., speech, music, and natural scenes. While auditory cortex (ACx) is thought to contribute to such processing, the underlying cortical mechanisms remain poorly understood. In a recent study (1), we investigated the processing of dynamic stimuli in mouse ACx, quantifying neural discriminability of dynamic sounds in ACx, and found that optogenetic suppression of parvalbumin (PV) neurons degraded discrimination performance, and specifically temporal coding, in ACx. In this study, we present a network model to address how PV neurons shape cortical discrimination performance.

PV neurons in ACx have been found to respond to both onset and offset in sounds (2, 3), suggesting they are specialized for rapid encoding of stimulus boundaries (4). Additionally, previous models have shown that the short-term synaptic plasticity (STP) profiles of inhibitory interneurons are critical for temporal context-dependent modulation (5). Here, we implemented a multi-layer cortical network model with excitatory neurons, and PV neurons responding to both onset- and offsets to explain the experimental observations in (1). The model reveals that PV neurons, and synaptic depression from PV to excitatory synapses are critical in sculpting cortical responses to dynamic stimuli.

## Materials and Methods

### Circuit structure overview

The model network is composed of three layers: the first layer representing pre-cortical input (IC, input cell), and the second (intermediate) and third (output) layer both representing ACx. Both cortical layers consist of excitatory cells and inhibitory parvalbumin-expressing (PV) cells.

### Architecture

The input layer consists of one input channel, based on previous experimental and modeling studies (6, 7). While previous experimental studies consisted of four spatial locations, our focus in this study was to dissect the effects of “within-channel” inhibition-mediated by PV neurons. Thus, we simplified the model architecture to focus on a single channel, which contains two excitatory subchannels that respond to different temporal features, i.e., onsets and offsets, of auditory stimuli. In each layer, both excitatory subchannels converge on a parvalbumin neuron, which then inhibits the excitatory cells in the next layer. Our goals were to match the discrimination performance and firing rate of excitatory neurons recorded in mouse ACx (1).

### Stimuli

For simulations of responses from (8), target stimuli consisted of two white noise tokens modulated by human speech-shaped envelopes with a sampling frequency of 200 kHz.

### Input spike generation via spectro-temporal receptive fields

Input responses were generated from spectro-temporal receptive fields (STRFs). The STRFs for the input layer were generated using the STRFlab Matlab toolbox (9) and modeled as the product of independent spectral and temporal kernels:

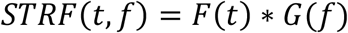

The temporal kernel was modeled as the difference of two linear functions (10, 11):

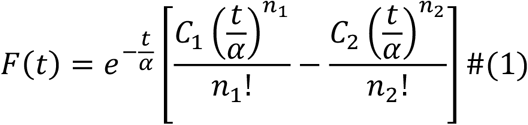

C_1_ and C_2_ control the weights of the excitatory and inhibitory regions of the temporal kernel, respectively. C_1_ was fixed at 1, while C_2_ was fixed at 0.88 to allow for a broader spiking profile.

α, which controls the timescale of the kernel, was fixed at 9.7 ms, while n_1_ was fixed at 5 and n_2_ was fixed at 8. Using this kernel allowed us to capture the temporal profile of STRFs measured in the mouse medial geniculate body (12).

The spectral kernel of the STRF was modeled as a 2D Gabor filter (13):

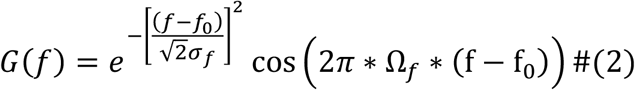

The frequency range is determined by the best frequency f_0_, the spectral bandwidth σ_f_, and modulation frequency Ω_f_. Respectively, these parameters were fixed at 4300 Hz, 2000 Hz, and 50 µs to generate a broadband STRF for all simulations.

The convolution of the STRF and stimulus spectrograms generated a firing rate profile, which was used to drive a Poisson spike generator. Refractoriness was modeled using a sigmoidal recovery function (14):

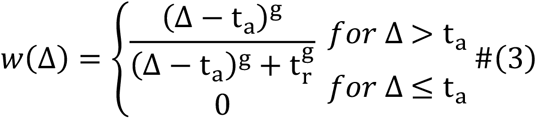

Absolute refractory period (τ_a_) was fixed at 1 ms, the relative refractory period (τ_r_) was fixed at 1 ms, and the timescale of the recovery function (γ) was fixed at 2 ms. For each of the two target stimuli, 10 trials were generated, each using a novel random token for Poisson spike generation.

To model responses to offsets, the convolution between the STRF and stimuli was inverted such that the minimum and maximum values were switched:

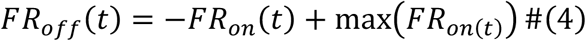

A similar scheme has been used in previous models to approximate offset responses in auditory cortex (15). To prevent evoked spiking at offset cells before stimulus playback, FR_off_ was set to zero for all times before its first zero-crossing. Finally, FR_off_ was thresholded such that all negative values were set to zero.

### Model structure and parameters

We utilized the DynaSim toolbox (16) to implement the model of mouse ACx. Neurons were modeled as leaky integrate-and-fire units with membrane voltage V:

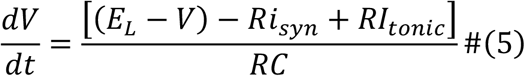

where E_L_ is the equilibrium potential, R is the membrane resistance, i_syn_ is the synaptic input current, I_tonic_ is applied tonic current, and C is the membrane capacitance. Universal parameters, such as capacitance and threshold voltage, are given in Table 1. Membrane resistances were varied between cell types to approximate the differences in integration time constants (Table 2), with PV cells having a faster membrane time constant than that of excitatory cells (5). Additionally, resting membrane potentials for PV cells were set higher than those of excitatory cells to ensure more rapid firing. When membrane voltage passed threshold, voltage was shunted to V_reset_ for an absolute refractory period t_ref_. In excitatory cells, spike-rate adaptation was also implemented (17):

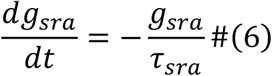

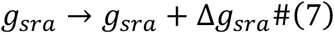

**Table 1.** Universal DynaSim parameters.

|  |  |
| --- | --- |
| C [nF] | 0.1 |
| $V_{\text{thresh}}$ [mV] | -47 |
| $I_{\text{tonic}}$ [nA] | 0 |
| Simulation time [ms] | 3500 |
| Integration time step [ms] | 0.1 |
| IC input gain | 0.165 |

**Table 2.** Unit parameters. DynaSim parameters that differed between units.

| Unit | R (MΩ) | $\Delta g_{\text{sra}}$ (nS/spk) | $\tau_{\text{sra}}$ (ms) | $t_{\text{ref}}$ (ms) | $E_L$ (mV) | $V_{\text{reset}}$ (mV) |
| --- | --- | --- | --- | --- | --- | --- |
| IC | 200 | 0 | -- | 1 | -65 | -54 |
| E | 200 | 0.3 | 100 | 1 | -65 | -54 |
| PV | 100 | 0 | -- | 0.5 | -57 | -52 |

After a post-synaptic spike, adaptation conductance increases by Δg_sra_ and recovers to zero with recovery time τ_sra_ (18). We found that implementing refractoriness and spike-rate adaptation in this manner allowed the model to match the firing rates modulations in experimental data.

To simulate network noise, the output excitatory cell received a Poisson-like noise input with a refractory period of 1 ms and a firing rate of 8 Hz, which matched the average spontaneous activity in experimental data. To decrease simulation time, network noise was not modeled using a separate unit; rather, tokens for Poisson spike times were generated before integration of cell voltage.

### Synaptic connections

Throughout simulations, certain synaptic weights were fixed. Within the circuit, the synaptic connectivity between excitatory cells (IC->E_Int_, E_Int_->E_Out_) was kept constant. Additionally, the synaptic conductance of the inputs to PV cells (IC->PV_Int_ and E_Int_->PV_Out_) was fixed to ensure that a PV neuron would fire around 3 ms before an excitatory cell receiving the same unitary input to approximate spike-time latencies observed in mouse auditory cortex (19). Additionally, all synapses featured short-term synaptic depression, based on previous models (20).

Post-synaptic currents (PSCs) were modeled using the difference of two decaying exponential functions with time constants τ_1_ and τ_2_, with τ_1_ > τ_2_:

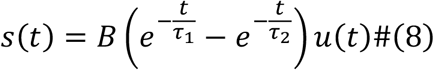

where B is a normalization constant such that s = 1 at maximum, and u(t) is the unit step. For simulations, s was rewritten as two ordinary differential equations:

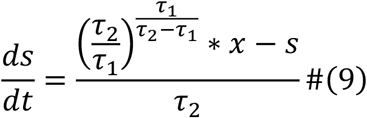

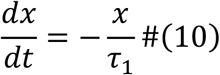

where x defines spike inputs to the post-synaptic cell. When a pre-synaptic spike is detected, x is increased by synaptic strength P. P is then decreased by a fraction (f_P_) of its current value (20):

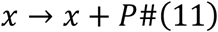

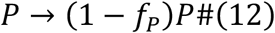

In the absence of pre-synaptic spikes, P recovers to 1 with time constant τ_P_:

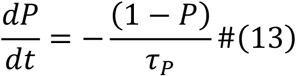

The full equation for post-synaptic currents is given as:

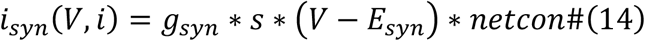

where g_syn_ is synaptic conductance; E_syn_ is reversal potential; and *netcon* is a binary matrix of network connections between cell populations, with rows representing sources and columns representing targets.

Values of τ_1_, τ_2_, and E_syn_ were based on recorded excitatory and inhibitory PSCs in mouse cortex (19) and kept constant throughout all simulations (21). Table 3 outlines the parameters for all synaptic connections.

**Table 3.** Synaptic parameters. DynaSim parameters that differed between synapses. N represents simulated network noise input from a Poisson unit in the output layer to model network noise in ACx.

|  |  |  |  | Nocon et al. (2023) |  |  |
| --- | --- | --- | --- | --- | --- | --- |
| Connection | $\tau_1$<br>(ms) | $\tau_2$<br>(ms) | $E_{\text{syn}}$<br>(mV) | $g_{\text{syn}}$<br>(nS) | $f_p$ | $\tau_p$ (ms) |
| E->E | 0.7 | 1.5 | 0 | 20 | 0.1 | 30 |
| N->E<sub>Out</sub> | 0.7 | 1.5 | 0 | 15 | 0 | -- |
| On->PV | 0.1 | 1 | 0 | 20 | 0.2 | 80 |
| Off->PV | 0.1 | 1 | 0 | 5 | 0 | 80 |
| PV->E<sub>On</sub> | 1 | 4.5 | -80 | 25 | 0.5 | 120 |
| PV->E<sub>Off</sub> |  |  |  |  |  |  |

### Biological rationale

STRFs at the IC were based on experimentally recorded data in mouse auditory neurons (12). Here, we start with the assumption that the model contains two excitatory subchannels that capture different features of auditory stimuli, with one responding to sound onsets and the other responding to sound offsets consistent with past studies that have reported parallel processing of onset and offset-responding cells in mouse auditory cortex (3). We model PV neurons as having both onset and offset components, the most frequently observed finding experimentally (2). Previous studies have shown the existence of both excitatory and inhibitory neurons within cortex (22–24), and other studies have found that PV cells encode gaps between bursts of white noise in auditory cortex (4).

### Calculating discrimination performance using spike distance measures

Neural discrimination performance was calculated using a template-matching approach from previous experimental and modeling studies (6, 7). Briefly, spike trains were classified to one of the two target stimuli based on whose template, one from each stimulus, yielded a smaller spike distance. For each simulation set of 20 trials, 100 iterations of template matching were done, in which one of the 10 spike trains for each target was chosen as a template, and all remaining trials were matched to each template to determine target identity. All possible pairs of templates were used across the 100 iterations to calculate an average value of performance. To ensure that discrimination performance was only based on spiking during stimulus playback, spiking between the simulation times of 300 ms and 3300 ms was used for analysis. This approach utilized spike train distances as inputs, and several measures were used, each of which capture different features of responses: SPIKE-distance; ISI-distance; rate-independent (RI) SPIKE-distance. Previous analysis also utilized spike count distance, the difference in number of spikes between trains, but discriminability based on this measure was close to chance for most data. Thus, spike count distance-based performance was excluded from this study.

SPIKE-distance (25, 26) calculates the dissimilarity between two spike trains based on differences in spike timing and instantaneous firing rate without additional parameters. For one spike train in a pair, the instantaneous spike timing difference at time t is:

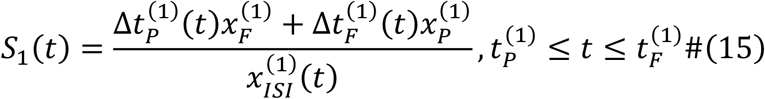

where Δt_P_ represents the distance between the preceding spike from train 1 (t_P_^(1)^) and the nearest spike from train 2, Δt_F_ represents the distance between the following spike from train 1 (t_F_^(1)^) and the nearest spike from train 2, x_F_ is the absolute difference between t and t_F_^(1)^, and x_P_ is the absolute difference between t and t_P_^(1)^. To calculate S_2_(t), the spike timing difference from the view of the other train, all spike times and ISIs are replaced with the relevant values in train 2. The pairwise instantaneous difference between the two trains is calculated as:

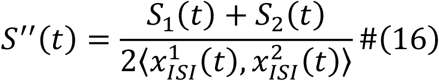

Finally, S_1_(t) and S_2_(t) are locally weighted by their instantaneous spike rates to account for differences in firing rate, and the total distance between the two spike trains is the integral of the profile:

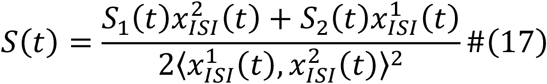

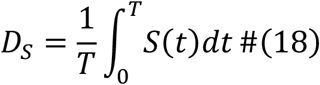

ISI-distance calculates the dissimilarity between two spike trains based on differences in instantaneous rate synchrony. For a given time point:

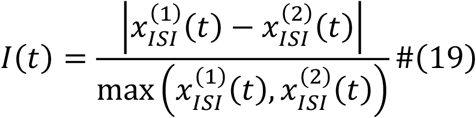

This profile is then integrated along the spike train length to give a distance value, with values of 0 obtained for either identical spike trains or pairs with the same constant firing rate and a global phase shift difference.

The RI-SPIKE-distance is rate-independent, as it does not take differences in local firing rate between the two spike trains into account. From SPIKE-distance calculations, the final step of weighing S_1_(t) and S_2_(t) by their instantaneous spike rates is skipped, yielding:

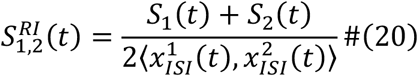

Like the other measures, the dissimilarity profile is integrated to give a distance value, with a value of 0 obtained for two identical spike trains. cSPIKE was used to calculate SPIKE-, ISI-, and RI-SPIKE distances between all possible spike train pairs (27).

### Parameter searches for synaptic depression strengths

To determine how the dynamics of synaptic depression affect spiking activity within the circuit, *f*_*P*_ was varied for each synapse type (E->E, PV->E, and E->PV). For all simulations, all three distance-based performance values at the output excitatory unit were calculated, along with the average firing rate during stimulus playback. The values of *f*_*P*_ that yielded firing and discriminability values approximating those seen in experimental data were then chosen for the rest of the simulations in this study.

### Optogenetic suppression of PV neurons

To replicate the experimental effect of optogenetic suppression of PV cells, all PV neurons received a hyperpolarizing current of 0.03 nA, and the mean firing rate of network noise inputs to the output excitatory unit was increased to 30 Hz to match the rise in spontaneous activity during suppression. The first effect was modeled to replicate the effect of increased spiking across all cells within experimentally recorded data, while the second effect captured the downstream effects of increased noise via network perturbation. To account for the variability from Poisson-like noise at output, we ran 30 simulations of the same set of 20 trials (10 per target stimulus) for each condition.

## Results

### Mouse ACx units exhibit high neural discrimination of temporally dynamic stimuli

To better understand cortical coding of complex scenes in a mouse model amenable to circuit manipulation using genetic tools, we recently utilized a cocktail party-like experimental paradigm (6) while recording from neurons in ACx (Figure 1A) (1). Specifically, we recorded responses to spatially distributed sound mixtures to determine how competing sound sources influence cortical coding of stimuli. The recording configuration consisted of 4 speakers arranged in horizontal space around the mouse at 4 locations on the azimuthal plane: directly in front (0º), two contralateral (90º and 45º) and one ipsilateral (−90º) to the right auditory cortex recording area. Two target stimuli were used, each consisting of a white noise token modulated by human speech-shaped envelopes from a speech corpus (28).

**Figure 1.**
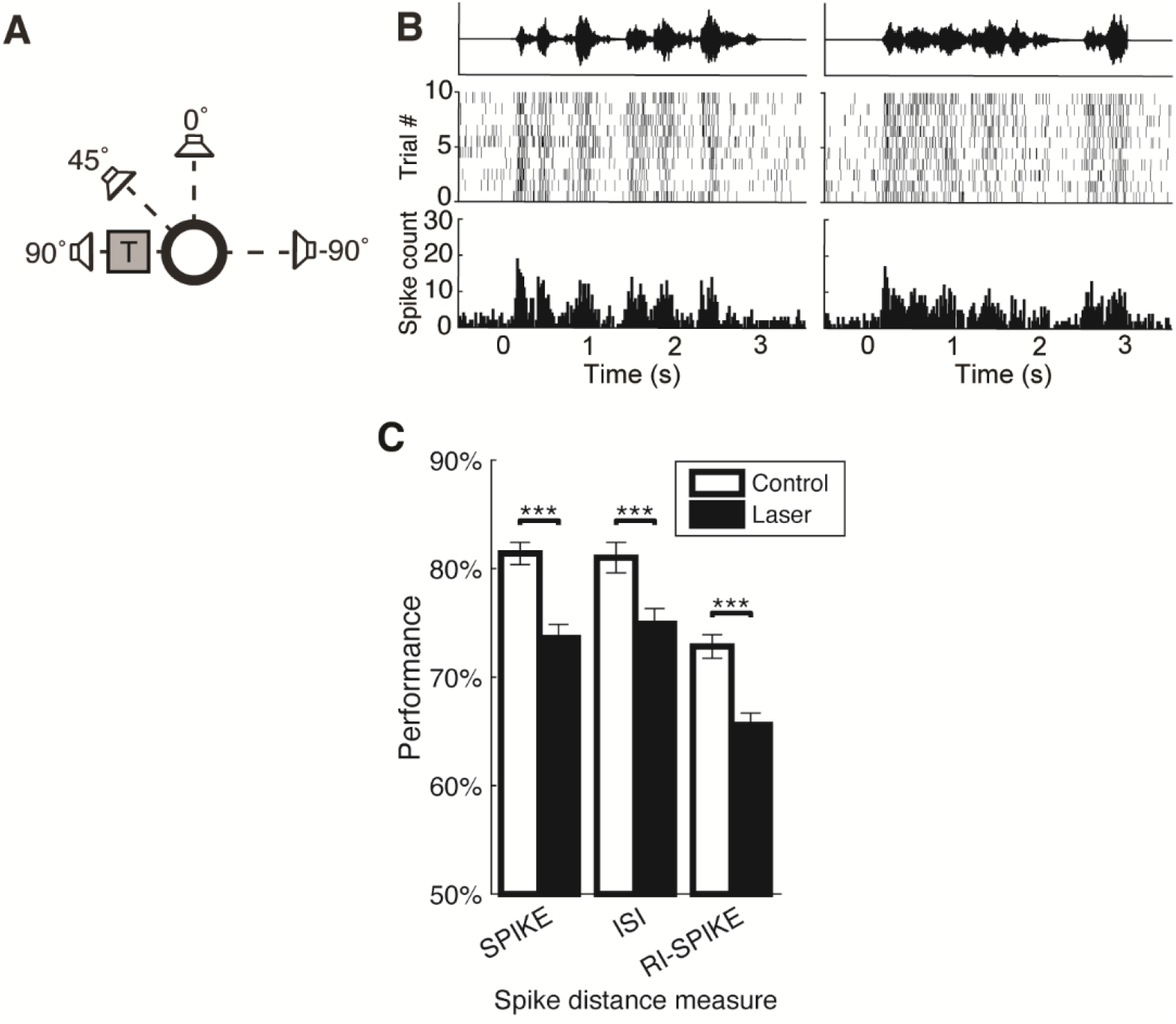
Experimental data from mouse ACx. **(A)** Diagram of experimental setup and spatial location of target stimuli. During trials, one of two target stimuli consisting of white noise modulated by a human speech-shaped envelope was played from one of the four speakers along the azimuth in horizontal space. **(B)** Responses to spatial configuration shown in A where the two targets originated at 90°, from an example single unit in mouse ACx. Responses followed the amplitude envelope of both targets, resulting in high SPIKE-distance-based neural discrimination performance (92%) at this configuration. **(C)** Average discrimination performance using different spike distance metrics across the population of 23 high-performing single units (*n* = 49 hotspots during clean trials). Paired t-tests found significant differences between control and optogenetic trials for all spike distance-based performances (SPIKE-distance: *p* = 8e-10; ISI-distance: *p* = 3e-08; RI-SPIKE-distance: *p* = 1e-08).

Figure 1B illustrates responses from an example single-unit cell during target-only conditions. Here, the spike rasters and peri-stimulus time histograms (PSTHs) follow the amplitude envelope of each target stimulus. To quantify the discriminability between the two target stimuli, we utilized a template matching-based classifier (see: Materials and Methods, ‘Calculating discrimination performance using spike distance measures’). The input to this classifier was one of several spike train distance measures: SPIKE-distance, which measures differences in spike timing and firing rate modulation between trains (27); inter-spike interval (ISI)-distance, which only measures differences in firing rate modulation; rate-independent (RI)-SPIKE-distance, which measures differences in spike timing while being independent of firing rate differences. We focused on a subset of single-unit neurons that exhibited high SPIKE-distance-based performance (above 70%) at one or more locations in the spatial grid that we refer to as hotspots. Across the population of hotspots, we found that SPIKE-distance-based performance was highest, followed by ISI-distance-based performance and then RI-SPIKE-distance-based performance (Figure 1C).

To investigate the role of PV interneurons in auditory discrimination performance, we compared discrimination performance at these hotspots with and without optogenetic suppression of PV neurons in ACx. Upon optogenetic suppression, we found that performance significantly decreased for all distance measures. These results indicate that optogenetic suppression disrupts both timing and rate-based coding within ACx.

We organized hotspots based on the order of SPIKE-distance-, ISI-distance, and RI-SPIKE-distance-based performances. In hotspots where firing rate-based (i.e., ISI-distance) performance was lowest (Figure 2A), we found that spike timing-based (i.e., RI-SPIKE-distance) performance primarily drove the high SPIKE-distance-based performance. When ACx was optogenetically suppressed, both the overall and spike timing-based performance was reduced, while firing rate-based performance remained unchanged. Hotspots where firing rate-based distance was second highest (Figure 2B) or highest (Figure 2C) indicate values where high SPIKE-distance-based performance was driven by firing rate modulations. Upon optogenetic suppression, all three performance measures were degraded at these hotspots. Across all three types of hotspots, we found those with higher rate-based performances tended to have higher firing rates during stimulus playback, which increased during optogenetic suppression (Figure 2D).

**Figure 2.**
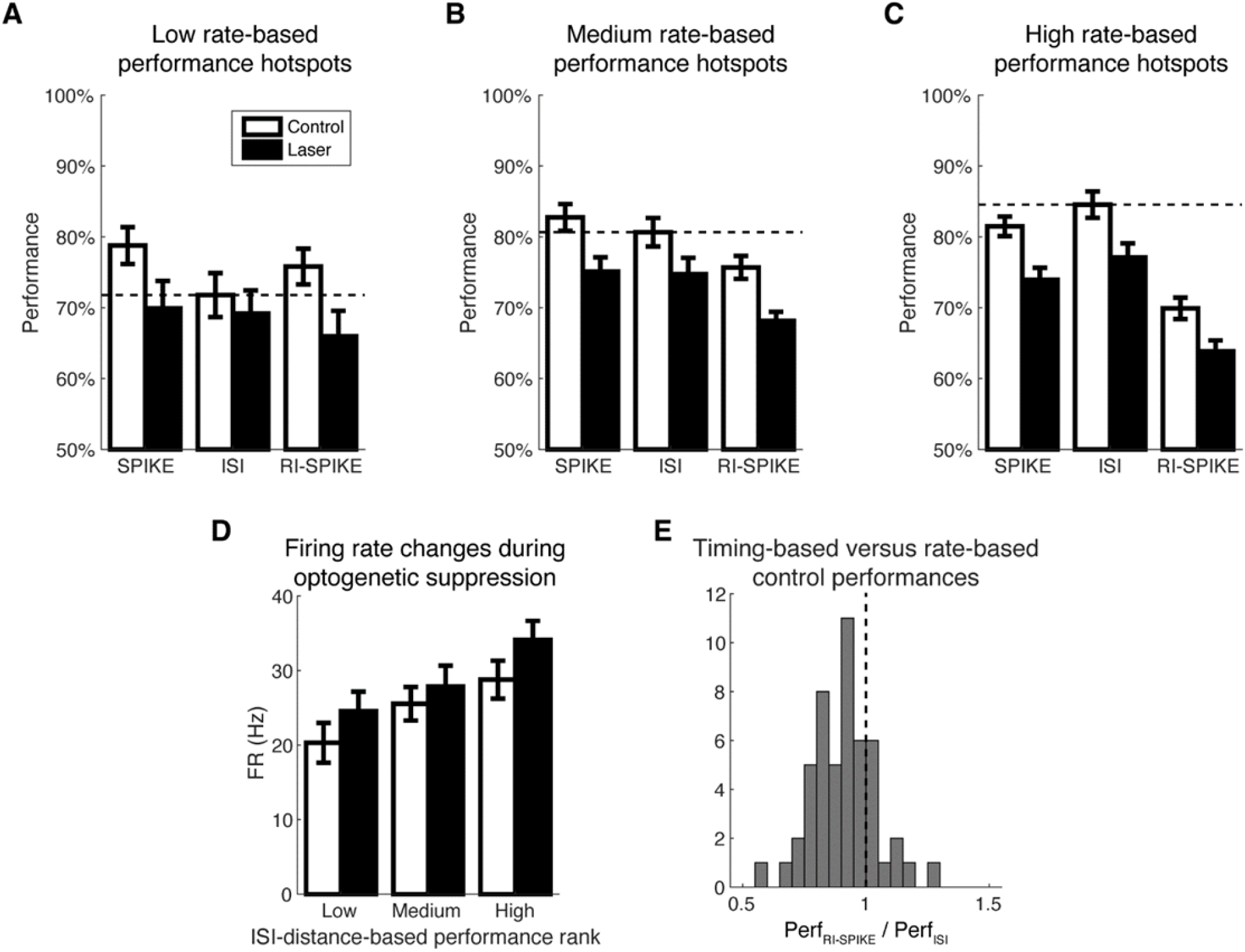
Hotspots in single-unit experimental data reflect a broad range of timing and rate-based contributions to performance. Hotspots in single-unit data were sorted by comparing firing rate modulation-based (ISI-distance) performance to the other two measures (SPIKE-distance, firing rate modulation and spike timing; RI-SPIKE-distance, spike timing only). **(A)** Hotspots where ISI-distance-based performance was the worst out of all three performance measures (n = 9), as indicated by the horizontal dashed line. **(B)** Hotspots where ISI-distance-based performance was higher than RI-SPIKE-distance-based performance but not SPIKE-distance-based performance (n = 15). **(C)** Hotspots where ISI-distance performance was higher than the other two performance values (n = 25). **(D)** Auditory stimulus evoked average firing rates during sound alone and PV optogenetic inhibition for each of the three hotspot types. (**E**)Histogram showing all 49 hotspots from single-units, sorted by the ratio between RI-SPIKE-distance-based performance and ISI-distance-based performance, with a ratio of 1 indicated by the dashed line. Ratios lower than 1 indicate hotspots where high neural discrimination performance is primarily driven by modulations in firing rate, rather than spike timing.

Because the average SPIKE-distance-based performance was similar across all types, we calculated the ratio of spike timing-based performance to firing rate-based performance for each hotspot (Figure 2E). This measure allowed us to determine the extent to which SPIKE-distance-based performance was based on either spike timing or firing rate modulation, with higher ratios emphasizing the former.

### Network Model: Distinct synaptic depression profiles across excitatory and PV inhibitory synapses support high discrimination performance

We created a multi-layer network model to replicate temporal responses to dynamic stimuli in mouse ACx neurons (Figure 3A). The input layer consists of an onset cell and offset cell with Poisson firing with absolute and relative refractory periods of 1ms. Within the cortical intermediate and output layers, the excitatory cell from the preceding layer synapses onto a feed-forward inhibitory circuit consisting of one excitatory cell (E) and one inhibitory PV cell (red) (Figure 3B). Table 1 shows the universal simulation parameters, while Table 2 shows the parameters for all cells in this model.

**Figure 3.**
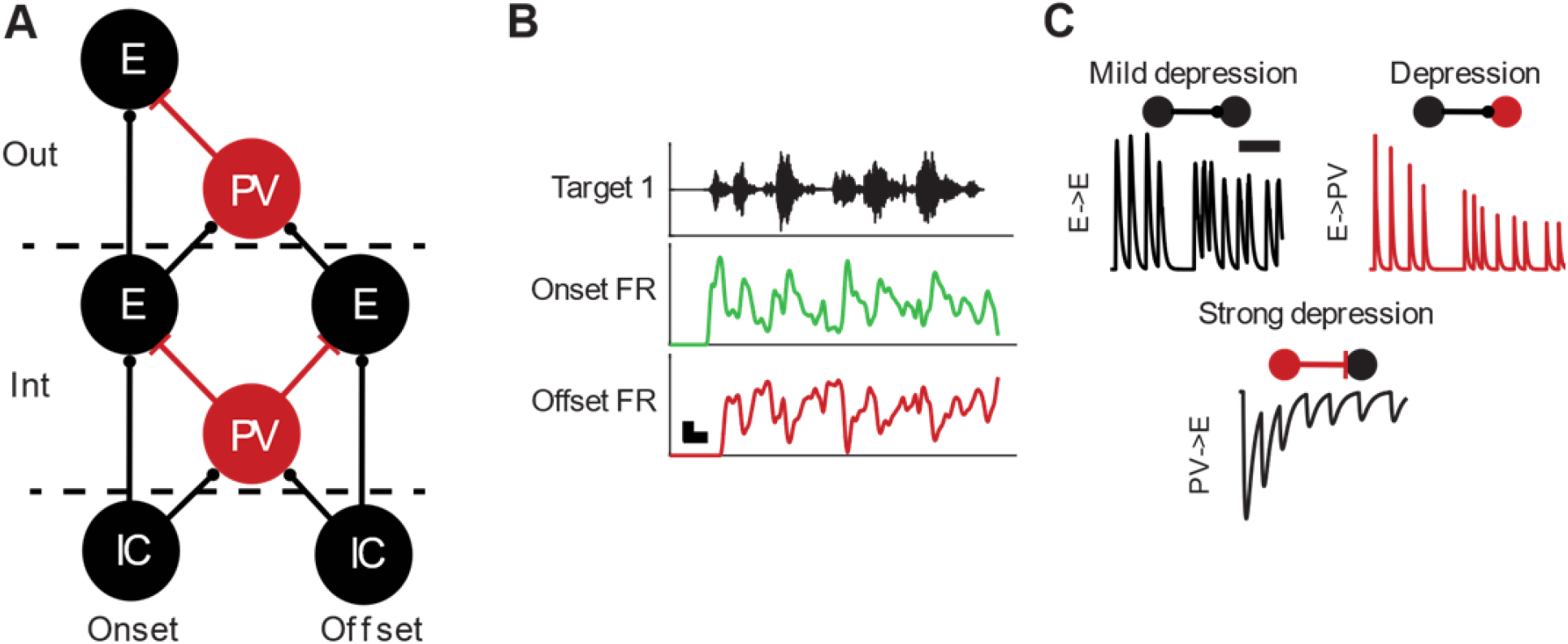
Computational model of auditory cortex layers and thalamic input. **(A)** Three-layer ACx model. The input stage of the model consists of a Poisson-like thalamic neuron with input from the inferior colliculus (IC) with both onset and offset type inputs. Firing rate of onset units is based on the convolution of an MGB STRF with stimulus spectrograms. Offset responses were generated by inverting the convolution between the STRF and stimulus and adding the absolute maximum value of the resulting firing rate trace. Both the intermediate (Int) and output (Out) layers exhibit feed-forward inhibitory circuits mediated by PV cells (red). PV neurons have both onset and offset components in their responses, the most frequent pattern observed experimentally (2). The excitatory cell in the output layer exhibited Poisson noise to simulate network noise in ACx. **(B)** Target 1 waveform with example FR trace from STRF convolution for onset-(red) and offset-responding (green) units. **(C)** Example traces of synaptic plasticity. Left: mild synaptic depression between excitatory cells. Right: synaptic depression in excitatory inputs to PV cells. Bottom: strong synaptic depression at inhibitory inputs to excitatory cells.

In all synapses, short-term plasticity (STP) was implemented as previously described (20). The model parameters are consistent with previous modeling studies in auditory cortex (5). Specifically, all synapses in this circuit featured depression, the strength of which was mildest between excitatory cells, followed by excitatory inputs to PV units, with the strongest depression in synapses from PV to excitatory units (Figure 3C). Recovery time constants for each synapse were chosen to match the firing rate modulations in experimental data. Table 3 shows the parameters for all synapses in this circuit.

To investigate how changes in depression strengths affected model responses, we varied the synaptic depression strength (f_P_) in each synapse type (E->E, E->PV, and PV->E), ranging from zero (f_P_ = 0) to maximal depression (f_P_ = 1). We first varied the depression strength between excitatory cells (Figure 4A). We found that weak synaptic depression was necessary for high firing rates and performance values at the output excitatory cell. When synaptic depression strength increased, performance dropped towards chance values and firing at output dropped to the network noise rate of 8 Hz.

**Figure 4.**
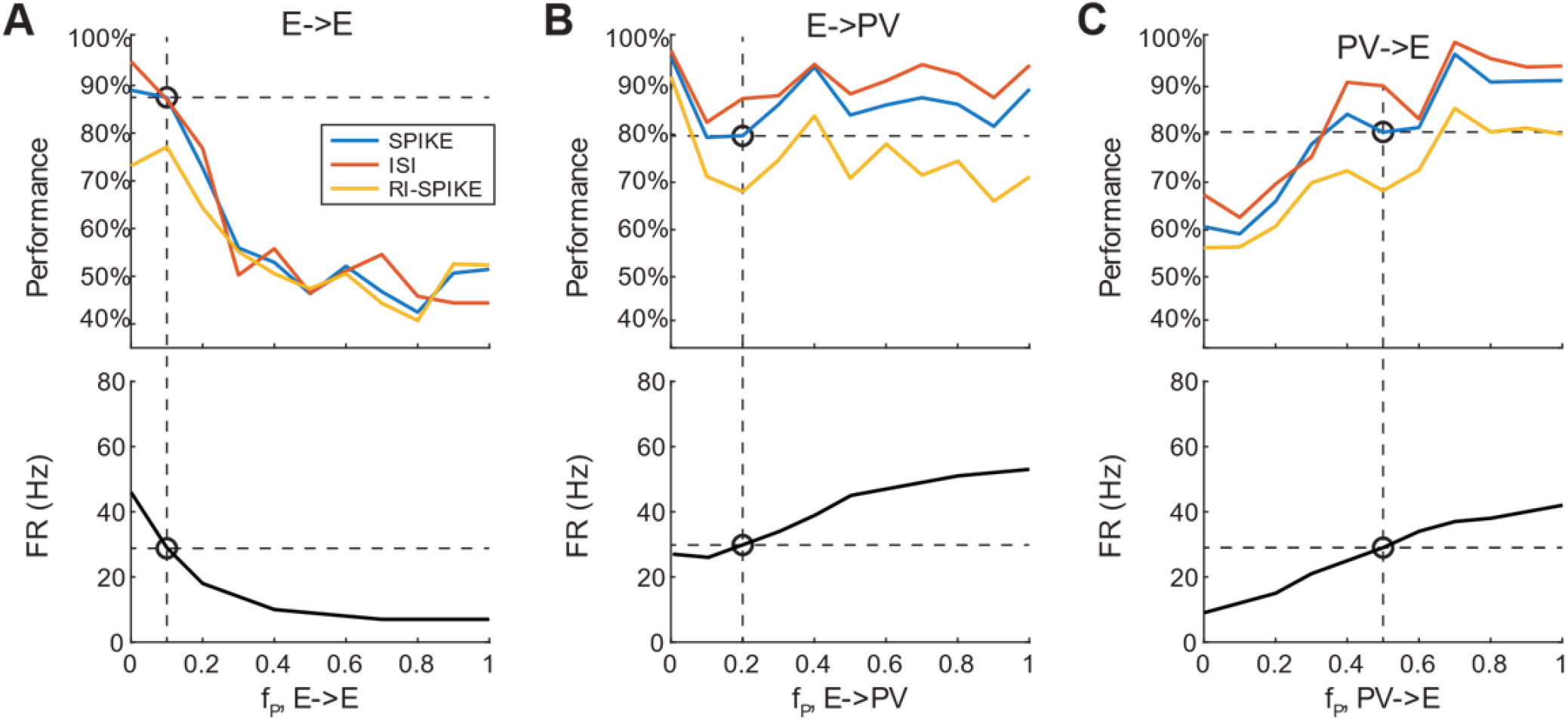
Effects of synaptic depression strengths on output discriminability and firing. Simulations for a rate-based hotspot in output excitatory cells, where synaptic depression strengths (f_P_) were varied while keeping all other parameters constant. **(A)** Synaptic depression strength between excitatory cells versus discriminability measures (top) and firing rate (bottom). Simulations where f_P,E->E_ **=** 0.1 (indicated by the vertical dashed line) leads to SPIKE-distance-based neural discriminability and firing rate values that approximate those of experimental data in hotspots with high firing rate-based performances, both of which are indicated by horizontal dashed lines in the upper and lower panels. **(B)** Synaptic depression strength between excitatory inputs to PV cells versus discriminability measures (top) and firing rate (bottom). Simulations where f_P,E->PV_ **=** 0.2 results in performance and firing rate values that approximate those of experimental data. **(C)** Synaptic depression strength of inhibitory inputs to excitatory cells versus discriminability measures and firing rate. Simulations where f_P,PV->E_ **=** 0.5 resulted in performance and firing rate values that approximate those in hotspots with high firing rate-based performance.

When we varied the synaptic depression strength at excitatory inputs to PV cells (Figure 4B), we found that all performance values did not greatly change at higher depression strengths, while firing rate greatly increased.

Finally, when we varied the synaptic depression strength of PV inhibition on excitatory cells (Figure 4C), we found that SPIKE-distance-based performance plateaued at higher values of f_P_, while ISI-distance-based performance increased, and RI-SPIKE-distance-based performance decreased. Additionally, firing rate steadily increased with synaptic depression strength. From these simulations, we chose depression strengths, i.e., f_P_ values that were consistent with the firing rates and performances that approximated the experimental single unit data from mouse ACx, as marked by the horizontal dashed lines in Figure 4. Thus, these results revealed that obtaining relatively high discrimination performance and firing rates consistent with the data requires relatively strong depression in PV->E synapses, moderate depression in E->PV synapses and mild depression in E->E synapses.

### Relative strengths of onset and offset component in PV neurons in model explains diversity of discrimination profiles observed experimentally

To model the diversity of rate-modulation hotspots (low, medium, and high) identified in the data, we next varied the strength between the onset and offset components preceding PV neurons and the PV neurons in both cortical layers while keeping all other circuit parameters constant. For hotspots with lower ratios of spike timing to firing rate-based performances, the offset component of PV neurons was reduced relative to the onset component. (Figure 5A).

**Figure 5.**
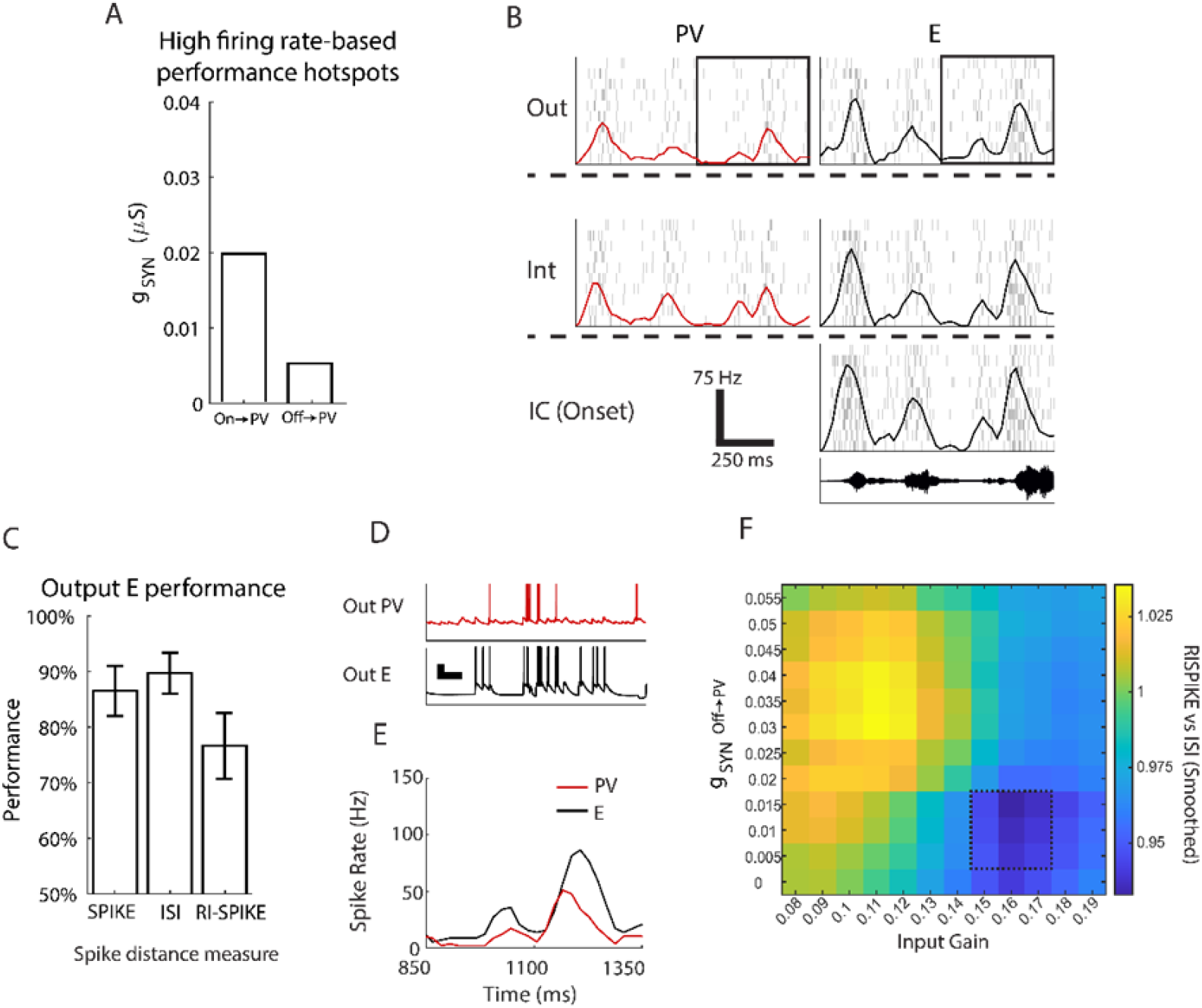
Hotspots with high firing rate-based performance can be modeled using PV neurons with a relatively high onset component, **(A)** To simulate hotspots where firing rate-based performance was higher than spike timing-based performance, inhibition strength from offset-sensitive PV units (On->PV, 20 µS) was reduced relative to that of onset-sensitive PV units (Off->PV, 5 µS). **(B)** Responses during the first second of one target stimulus (black waveform, bottom) from each cell, with rasters plotted in gray and the corresponding PSTH overlaid in the same colors as those shown in Figure 3A. To better show firing rate patterns in all cells, PSTHs were smoothed using a moving-average filter spanning 3 samples. Scale bars at bottom-right corner indicates vertical scale of PSTHs and horizontal time scale. Average firing rate at the output excitatory cell was 32 Hz during stimulus playback. **(C)** Performance measures of simulations shown in A. Here, ISI-distance-based performance is higher than RI-SPIKE-distance-based performance, indicating that the high SPIKE-distance-based performance is dominated by rapid changes in firing rate modulation. **(D)** Example voltage traces from output layer neurons. Onset-sensitive PV neurons (top) fire first from IC inputs, which then leads to inhibition of E cells (middle), thereby sculpting the ascending edge of firing rate modulations in excitatory cells. The descending edge is sculpted by offset-sensitive PV cells (bottom). Scale bars measure 20mV and 10ms. **(E)** Zoomed-in smoothed PSTHs (bin size 20ms) of output layer cells during the first half second of sound onset. PSTHs of PV cells indicate how inhibition shapes the firing rate patterns in excitatory cells. **(F)** Example parameter search showing approximate RI-SPIKE/ISI performance values testing IC input gain vs (IC_Off_->PV)g_syn_. Outlined section highlights ISI dominant region in parameter space (black outline). The color plot shows 7 by 7 zero-padded averaging convolution of a single parameter sweep (average of 49 cells per cell shown). Values were first subtracted by the grid mean and then the mean was reintroduced after convolution to reduce noise from zero padding.

Figure 5B illustrates the model’s response to one of the two target stimuli shown in Figure 1B. Throughout the ascending circuit, the firing rate pattern from each input is maintained. In the model, output units exhibit periods of firing rate modulation where spiking is close to zero during the quiet periods of the target stimulus.

We also found that this circuit effectively captures the pattern of firing rate-based versus spike timing-based performance in high rate-based performance hotspots observed experimentally (Figure 5C). For the simulations shown in 5B, ISI-distance-based performance was higher than SPIKE-distance- and RI-SPIKE-distance-based performance was consistent with the patterns observed in the high rate-based performance hotspots in the data (Figure 2C).

The model also captures the relative spike latencies of each cell type within cortex (Figure 5D). Given the same input from IC, the onset component of PV neurons fires first, which inhibits the excitatory PSC from the same spike input at E. These results are consistent with the differences in receptive fields between excitatory and PV neurons in mouse ACx (19). Due to synaptic depression, later spikes from the PV cell have a much weaker inhibitory effect on the excitatory cell, thus allowing the E cell to fire during this period of weakened inhibition. The descending edge of excitatory firing is then shaped by inhibition from the offset component in PV neurons, which sharpens firing rate modulations (Figure 5E).

The parameter space underlying the relative offset to PV strength in neurons versus input gain (Figure 5F), shows the distinction between the timing-based regime and the rate-based regime. The graph displays the approximate ratio between the timing-based regime and the rate-based regime (RI-SPIKE performance/ISI performance). A relatively high input gain and low offset component results in a rate-modulation dominated regime of coding (blue region).

To capture hotspots with higher ratios of spike timing to firing rate-based performances, we ran simulations where the offset component of PV neurons was higher than the onset component (Figure 6A). Within experimental data, we found that these hotspots exhibited responses where the noise floor firing rate during stimulus playback was lower than the spontaneous firing rate. Thus, we hypothesized that increased inhibition during offset periods facilitates higher spike timing-based performance. Figure 6B shows the responses to one of the two target stimuli across all cells within the circuit. The output excitatory cell exhibited an average firing rate of 20 Hz during stimulus playback, which approximates values seen in mouse ACx. For these simulations, SPIKE-distance- and RI-SPIKE-distance-based performances were almost equal, with ISI-distance-based performance being the lowest (Figure 6C), which agrees with the trends observed in the low rate-based performance hotspots in the data (Figure 2A). Voltage traces and firing rate modulation patterns at output neurons retain the same sharpening mechanism seen in simulations of hotspots with high firing rate-based performances (Figures 6D-E). Figure 6F illustrates the range of parameter values for the spike timing-dominated regime (highlighted region, 0.035 µS (IC_off_->PV)_gsyn_, 0.1 input gain), with relatively low input gain and a relatively high offset component. Thus, the model reflects the diversity of firing rate-based performance seen in experimental data and reveals that altering the relative onset and offset components of PV neurons effectively captures the broad range of coding profiles seen in mouse ACx.

**Figure 6.**
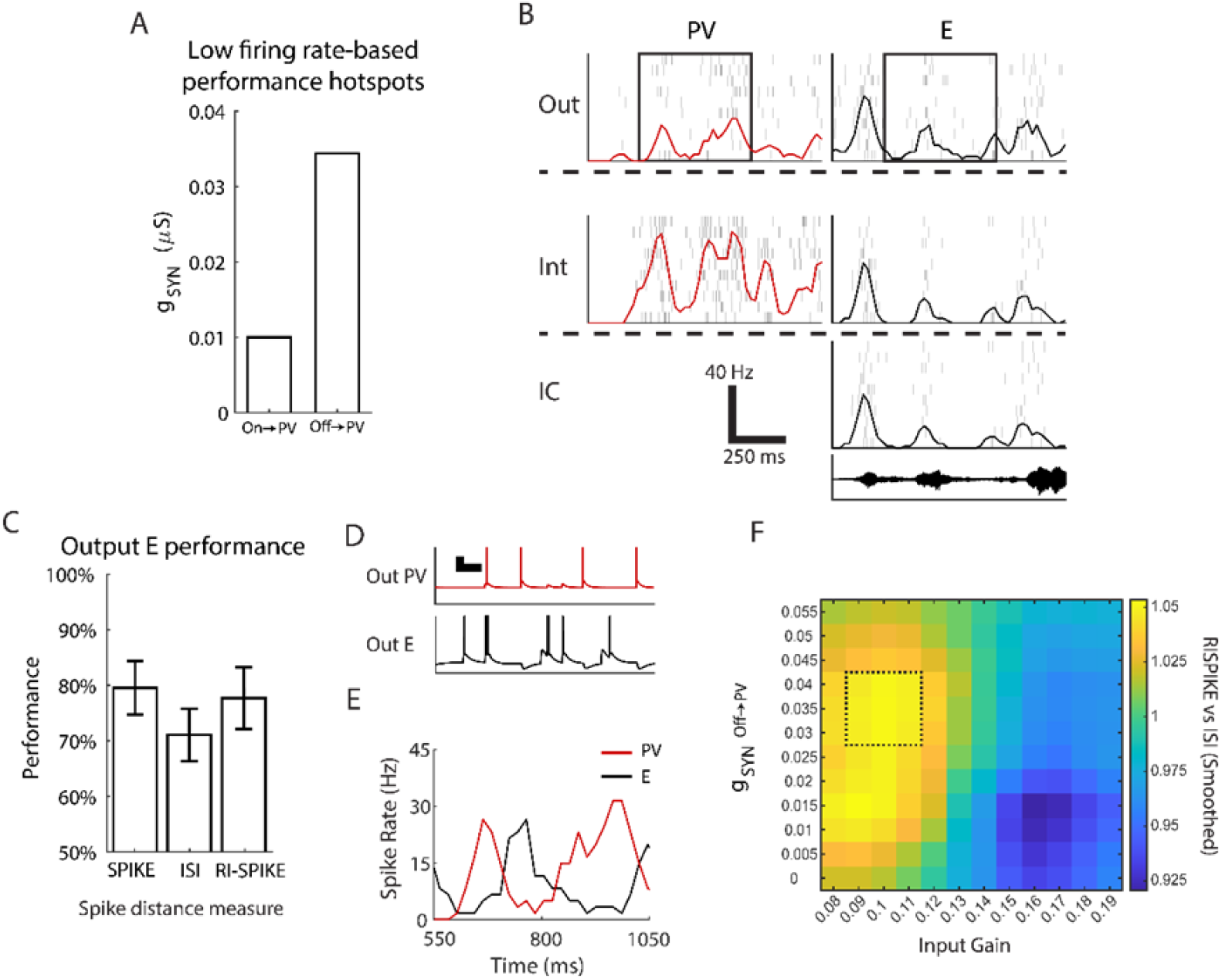
Hotspots with high spike timing-based performance can be modeled using PV neurons with a relatively high offset component. **(A)** To simulate hotspots where spike timing-based performance was higher than firing rate-based performance, inhibition strength from offset-sensitive PV units (w_Off_, green) was elevated relative to onset-sensitive PV units (w_On_, red). **(B)** Responses during the first second of one target stimulus (black waveform, bottom) from each cell, with rasters plotted in gray and the corresponding smoothed PSTH overlaid in the same colors as those shown in Figure 3A. Scale bars at bottom-right corner indicates vertical scale of PSTHs and horizontal time scale. Average firing rate at the output excitatory cell was 20 Hz during stimulus playback. **(C)** Performance measures of simulations shown in A. Here, ISI-distance-based performance is lower than RI-SPIKE-distance-based performance, indicating that the high SPIKE-distance-based performance is dominated by spike timing comparisons between trains. **(D)** Example voltage traces from output layer neurons. Onset-sensitive PV neurons (top) fire first from IC inputs, which then leads to inhibition of E cells (middle), thereby sculpting the ascending edge of firing rate modulations in excitatory cells. The descending edge is sculpted by offset-sensitive PV cells (bottom). Scale bars measure 20mV and 10ms. **(E)** Zoomed-in smoothed PSTHs (bin size 20ms) of output layer cells during the first half second of sound onset. (**F**)E Example parameter search showing approximate RI-SPIKE/ISI performance values testing IC input gain vs (IC_Off_->PV)g_syn_. Outlined section highlights RI-SPIKE dominant region in parameter space (black outline). Convolution method is the same as described in Figure 5F.

### Model explains experimentally observed effects of optogenetic suppression of PV neurons

We next employed the model to simulate the effects of reduced PV signaling and compared the results to empirical data collected in mouse ACx during optogenetic suppression of PV cells. For the optogenetic condition, we inhibited PV neurons in all layers with a hyperpolarizing current to model the elevated spontaneous firing rate observed in single unit recordings (1), we increased the mean firing rate of the Poisson-like network noise to simulate the increased excitability in the cortical network (Figure 7A).

**Figure 7.**
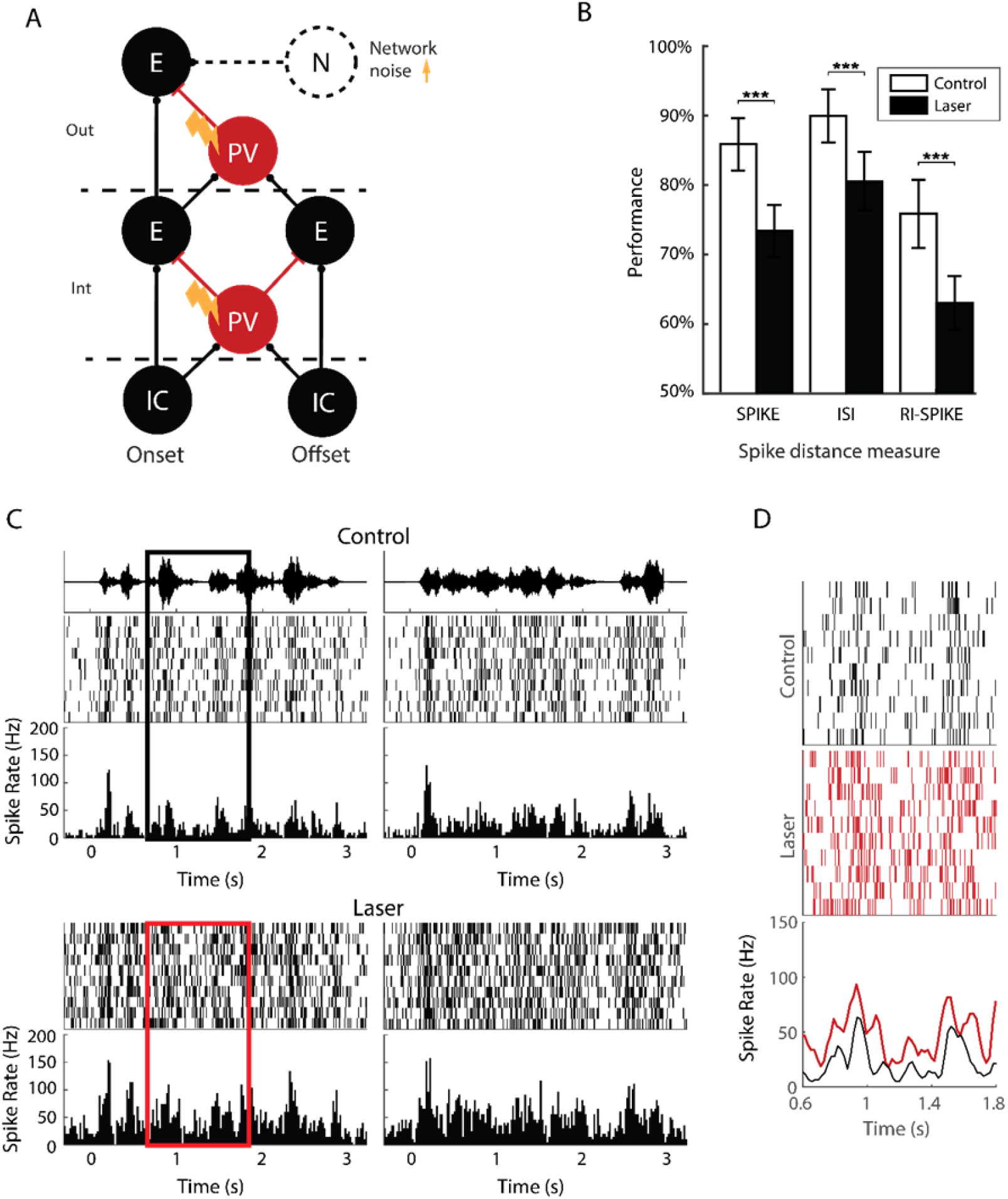
Simulations of responses during optogenetic suppression for a hotspot with high firing rate-based performance. For all simulations used in this figure, synaptic connectivity strengths were the same as in Figure 5. **(A)** Diagram showing changes to circuit during optogenetic suppression simulations. All PV cells are suppressed with a hyperpolarizing current of 0.03 nA, and the network noise input to the output excitatory cell is increased from 8 Hz to 30 Hz. **(B)** Average performance values during control and optogenetic conditions across 30 sets of simulations per condition, with error bars representing ± 1 standard deviation. The significance between laser and control trials for all distance metrics (p < 0.001) was calculated using Wilcoxon rank sum test. **(C)** Example rasters and PSTHs to each target stimulus during one control and one optogenetic simulation. **(D)** Inset showing zoomed-in portion of the response to target stimulus #1, between 0.6 and 1.8s after sound onset, as outlined in C. Responses during optogenetic trials show a higher noise floor, compared to the control condition.

In agreement with experimental results, we found that discrimination performance values at the output unit decreased with optogenetic suppression for all spike distance measures (Figure 7B). Control simulations were found to approximate the firing rate profiles found within mouse ACx, where the firing rate profile followed the temporal envelope of both target stimuli and showed very little firing during the quiet periods of both targets (Figure 7C, top). During simulations where PV neurons were suppressed, we saw a decrease in firing in all PV cells and a slight increase in firing at both cortical excitatory units (Figure 7C, bottom). Notably, we found that the noise floor was larger in optogenetic suppression trials than in the control condition during stimulus playback (Figure 7D).

When we simulated optogenetic suppression for low firing rate populations that show enhanced spike timing-based performance, the model replicated the experimental results across all three measures of temporal and rate dependent discriminability. Specifically, optogenetic suppression led to a decrease in both SPIKE-distance- and RI-SPIKE-distance-based performances, while ISI-distance-based performance remained unchanged (Figure 8A), which agrees with the data in Figure 2A. Like the simulations from the firing rate-based hotspot, responses during the control condition were found to approximate those within mouse ACx (Figure 8B, top). During optogenetic simulations, we saw the same increase in the noise floor during stimulus playback as in the rate-based hotspot simulations (Figure 8B, bottom, and Figure 8C). Thus, the three-layer circuit model captures the experimentally observed reduction in neural discriminability with optogenetic suppression of PV neurons in mouse ACx.

**Figure 8.**
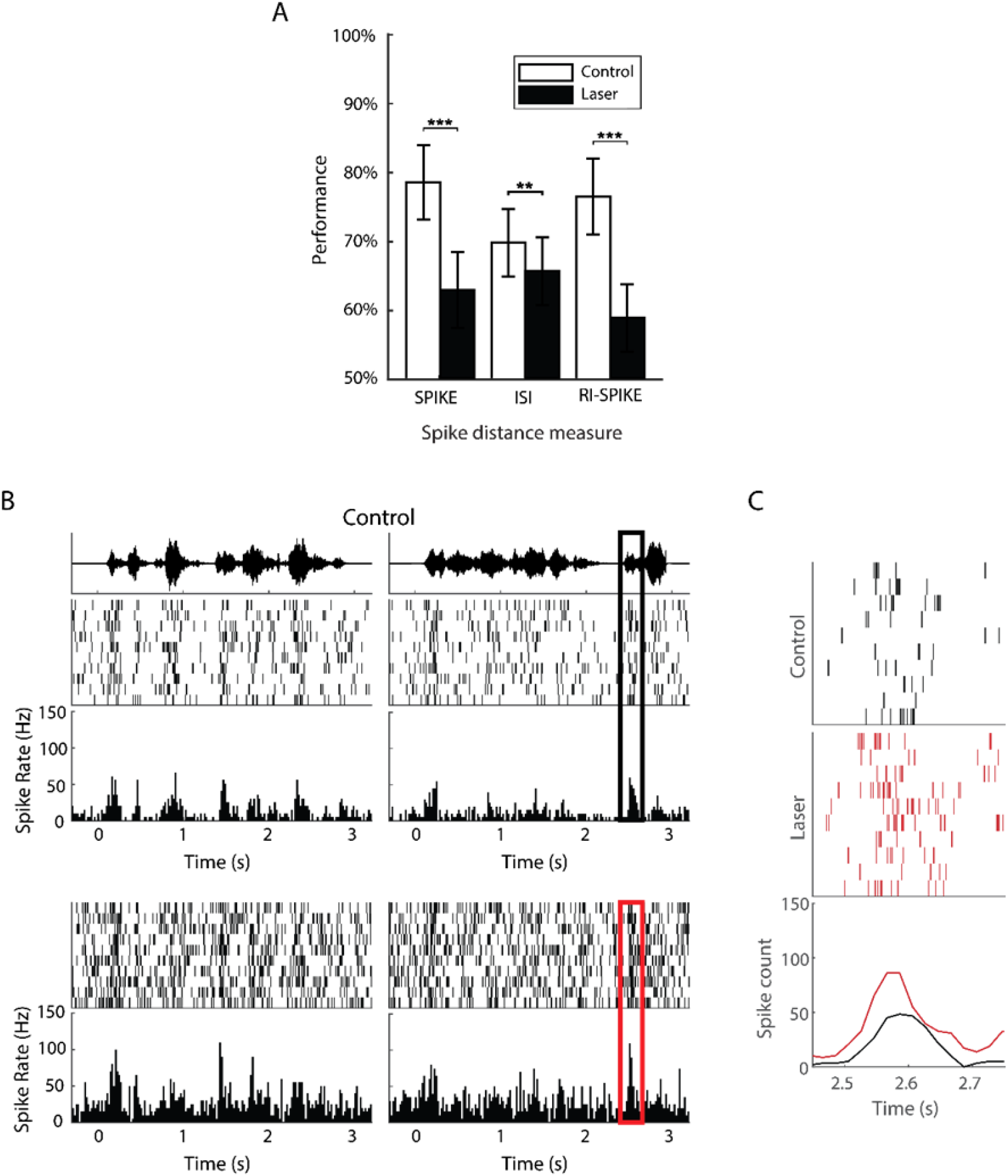
Simulations of responses during optogenetic suppression in a hotspot with high spike timing-based performance. For all simulations used in this Figure, synaptic connectivity strengths were the same as in Figure 6. **(A)** Average performance values during control and optogenetic trials across 30 sets of simulations per condition, with error bars representing ± 1 standard deviation. The significance between laser and control trials for SPIKE and RI-SPIKE distance (p < 0.001) and ISI distance (p < 0.01) was calculated using Wilcoxon rank sum test. **(B)** Example rasters and PSTHs to each target stimulus during one control and one optogenetic simulation. **(C)** Inset showing zoomed-in portion of the response to target stimulus #2, between 2.45s and 2.75s after sound onset, as outlined in B. Responses during optogenetic trials show a higher noise floor, compared to the control condition.

## Discussion

### Diverse neural discrimination performance profiles in ACx

In previous experimental study examining the effects of optogenetically suppressing PV cells in mouse ACx, we found that performance was degraded in terms of both spike timing, firing rate modulations, and a combination of the two (1). In this study we further analyzed the degree to which high neural discriminability was determined by either spike timing or firing rate modulations within each hotspot. We then set out to determine if we could construct a model that effectively captures the diversity of rate and temporal coding components seen in single unit data. In addition, we focused on revealing the properties by which PV neurons could influence discriminability across a range of hotspots present in mouse ACx.

### Network model explains diversity of profiles

Previous computational studies of the mouse auditory cortex primarily focused on modeling responses to transient stimuli such as tones and white noise bursts (5, 29). Here, we sought to replicate responses to longer, and temporally dynamic stimuli in mouse ACx. using experimentally recorded spectral temporal receptive fields in mouse MGB (12). This input is then temporally sharpened by intra-cortical processing via inhibition from PV interneurons.

One critical finding of the model is that the profile of synaptic depression parameters in distinct synapses is critical for attaining high neural discriminability in auditory cortex. Similarly, synaptic depression at inputs to PV cells and their inhibitory effect on excitatory cells were both found to facilitate rapid modulations in firing rate within the circuit. Specifically, as the thalamic input was processed in cortical layers, firing rate peaks were maintained, and offset components of PV neurons enhanced suppression of the troughs in firing rate, thereby enhancing firing rate modulations.

### Potential role of onset and offset components of PV interneurons in generating diverse discrimination performance profiles and responses to stimulus envelopes

We demonstrated that the model circuit was able to capture the diversity of discriminability profiles within auditory cortex by changing the relative strengths of onset and offset components of PV neurons. When onset components were relatively high compared to offset, excitatory units exhibited more prominent rate modulation dominated coding, i.e., low spike timing to firing rate-based performance ratios. When offset components of PV neural responses were relatively stronger compared to onset, the model captured the remaining set of hotspots with spike timing dominated coding (high spike timing to firing rate-based performance ratios). Previous studies have found that PV neurons are implicated in encoding gaps in noise within auditory cortex which suggests an important contributions to spike timing (4). A critical feature in our model is that PV neurons have both onset and offset components, based on experimental observations from (2).These components may arise from distinct parallel onset and offset inputs originating in the thalamus (3). Our model further elucidates the contribution of offset component in PV neurons in enhancing temporal coding of dynamic stimuli.

### Model limitations

The model in this study is a simplified version of cortical circuits, as it only includes feed-forward circuits between excitatory and PV units. It does not account for any recurrent inhibitory mechanisms between the two neuron types, nor does it include any other interneuron type, such as somatostatin-expressing (SOM) or vasoactive intestinal peptide-expressing (VIP) neurons, which might further facilitate the high neural discriminability seen in cortex. Additionally, the model utilized single synapses over a population model in which multiple inputs converge onto neurons. These simplifications allowed us to better dissect which components of the cortical network are integral for capturing the variety of hotspots seen in experimental data. The current results suggest an important role of onset and offset components in PV neurons to temporally sharpen cortical responses and improve cortical discrimination performance. Future models should incorporate other interneuron types to capture additional aspects of cortical processing. For example, SOM neurons may be important to capture “cross-channel” surround inhibition, and VIP neurons may be important for capturing top-down effects on cortical processing (30).

In the model, the responses of offset neurons were modeled phenomenologically similar to a previous study on auditory thalamic responses in mice (15). We found that this approximation produced responses like the responses of offset neurons in the data. Other studies have hypothesized that responses to stimulus offsets along auditory pathways are mediated by hyperpolarization-activated current, amplified by downstream mechanisms, or inherited from upstream regions (31, 32). Future studies should explore these mechanistic models of offset-responding neurons.

Finally, while the model utilizes STP between all synapses, the parameters chosen to model the temporal profile and strength of STP were selected to match the rapid firing rate modulations noted in the experimental data. The recovery times and depression strengths that were chosen for the circuit approximate the relative strengths of synaptic depression found within cortex, with mild depression between excitatory inputs to relay cells and strong depression in inhibitory inputs to excitatory cells (5). With these parameters, the model captured both the neural discrimination performance profiles and mean firing rate values seen within the experimental data.

### Key model predictions and future directions

In this study, we presented a computational model describing how PV-driven inhibition transforms thalamic inputs across cortical layers to facilitates high neural discrimination performance within auditory cortex. The model is based on physiological data in mice and posits that 1) cortical processing refines stimulus encoding at the thalamic level as spiking propagates downstream, 2) spiking in PV neurons are more phasic in time when compared to responses in excitatory neurons, 3) PV neurons with both onset and offset components sharpen spiking activity in excitatory neurons, thus leading to increased neural discriminability performance within cortex.

In this model, we focused on dissecting the rich temporal dynamics of “within-channel” inhibition mediated by PV neurons, to replicate experimental data from clean trials in which only a single stimulus was present. We did not model “cross-channel” interactions across spatial channels, which may be critical for modeling the emergence of hotspots at different spatial configurations. Based on the architecture of previous models (7, 30), future models of mouse ACx should incorporate multiple spatial channels, with cross channel “surround” inhibition. The analysis found that suppression of PV neurons reduced performance across all configurations of target and masker location (1); thus, we predict that other interneurons types, e.g., SOM and VIP neurons, may play an important role in determining the emergence of hotspots at different spatial configurations.

More generally, the modeling paradigm developed in this study may be useful in investigating the roles of different cortical interneuron types in complex scene analysis. The experimental data that were used to create and validate this model involved the optogenetic suppression of PV interneurons within mouse ACx. Future experiments with optogenetic manipulations or opto-tagging of other interneuron types within cortex, such as SOM or VIP neurons can be used to expand the model circuit. Ultimately, an understanding of the contributions of the diverse populations of interneurons in cortex, will be necessary to reveal cortical circuit mechanisms underlying complex scene analysis.

## Acknowledgements

We thank Michael Economo, Alberto Cruz-Martín, Monty Escabí, and Kristina Penikis for helpful discussions and comments on the manuscript.

